# LD Hub: a centralized database and web interface to perform LD score regression that maximizes the potential of summary level GWAS data for SNP heritability and genetic correlation analysis

**DOI:** 10.1101/051094

**Authors:** Jie Zheng, A. Mesut Erzurumluoglu, Benjamin L. Elsworth, Laurence Howe, Philip C. Haycock, Gibran Hemani, Katherine Tansey, Charles Laurin, Early Genetics and Lifecourse Epidemiology (EAGLE) Eczema Consortium, Beate St. Pourcain, Nicole M. Warrington, Hilary K. Finucane, Alkes L. Price, Brendan K. Bulik-Sullivan, Verneri Anttila, Lavinia Paternoster, Tom R. Gaunt, David M. Evans, Benjamin M. Neale

**Author notes:** To whom correspondence should be addressed. [† Joint senior authors].

## Abstract

**Motivation:** LD score regression is a reliable and efficient method of using genome-wide association study (GWAS) summary-level results data to estimate the SNP heritability of complex traits and diseases, partition this heritability into functional categories, and estimate the genetic correlation between different phenotypes. Because the method relies on summary level results data, LD score regression is computationally tractable even for very large sample sizes. However, publicly available GWAS summary-level data are typically stored in different databases and have different formats, making it difficult to apply LD score regression to estimate genetic correlations across many different traits simultaneously.

**Results:** In this manuscript, we describe LD Hub – a centralized database of summary-level GWAS results for 177 diseases/traits from different publicly available resources/consortia and a web interface that automates the LD score regression analysis pipeline. To demonstrate functionality and validate our software, we replicated previously reported LD score regression analyses of 49 traits/diseases using LD Hub; and estimated SNP heritability and the genetic correlation across the different phenotypes. We also present new results obtained by uploading a recent atopic dermatitis GWAS meta-analysis to examine the genetic correlation between the condition and other potentially related traits. In response to the growing availability of publicly accessible GWAS summary-level results data, our database and the accompanying web interface will ensure maximal uptake of the LD score regression methodology, provide a useful database for the public dissemination of GWAS results, and provide a method for easily screening hundreds of traits for overlapping genetic aetiologies.

**Availability and implementation:** The web interface and instructions for using LD Hub are available at http://ldsc.broadinstitute.org/

## 1 Introduction

There is now substantial empirical evidence demonstrating that the majority of complex traits and diseases in humans are influenced by hundreds if not thousands of genetic loci of small effect scattered across the genome as was first predicted a century ago (East 1916; Fisher 1918). The advent of high throughput micro-array genotyping and now next generation sequencing technologies has meant that genome-wide data can be leveraged to ask fundamental questions concerning the underlying genetic architecture of common complex traits and diseases including the degree to which genetic variation affecting complex phenotypes is tagged by SNPs on genome-wide arrays (Yang et al, 2010; Yang et al, 2011; Lee et al, 2012), the degree to which this variation represents different functional categories and/or biological pathways (Yang et al. 2011; Gusev et al. 2014; Finucane et al, 2015), and the extent to which genetic aetiologies are shared across different phenotypes (Lee et al. 2012; Lee et al. 2013; Bulik-Sullivan et al, 2015b). To date most of these types of analyses have been performed using genetic restricted maximum likelihood analysis (GREML) as implemented in software packages such as GCTA and LDAK (Yang et al, 2010; Yang et al, 2011; Lee et al, 2012; Speed et al. 2012). However these methods require individual-level genotype data, which is often not available as most of the largest GWAS analyses are conducted through meta-analyses, and so typically only report summary results statistics (Zheng et al, 2013). Additionally GREML can be computationally prohibitive when analyzing raw genome-wide SNP data from hundreds of thousands of individuals. Consequently, most GREML analyses reported in the literature to date have been hypothesis driven studies that have involved only a small number of related traits (Table 1).

**Table 1.** Comparison between GREML and LD Score Regression via LD Hub.

| GREML | LD Score regression via LD Hub |
| --- | --- |
| One dataset at a time | Integrates multiple GWAS results datasets |
| Run time depends on number of individuals and traits | Run time depends on number of traits only |
| Manual implementation | Automated |
| Usually one or a few traits at a time | Many traits simultaneously |
| Typically hypothesis driven | Hypothesis driven or hypothesis free |
| Computationally prohibitive for large numbers of individuals | Handles large numbers of individuals easily |

In order to address these limitations, Bulik-Sullivan et al previously proposed a different method, LD score regression (Bulik-Sullivan et al, 2015a). Essentially the method involves regressing summary results statistics from millions of genetic variants across the genome on a measure of each variant’s ability to tag other variants locally (i.e. its “LD score”). The intuition behind the approach is that if a trait is genetically influenced, then variants that tag more of the genome (i.e. have high LD scores) should have a greater opportunity to tag causal variants and therefore have higher test statistics on average than variants that have low LD scores. In this way genome-wide inflation of test statistics due to genuine polygenicity can be distinguished from biases such as population stratification and cryptic relatedness. The basic method is very flexible and can be adapted to estimate SNP heritability, calculate a more accurate and efficient genome-wide inflation correction factor than genomic control (Bulik-Sullivan et al, 2015a), partition the SNP heritability by functional category (Finucane et al, 2015), and estimate the genetic correlation between different complex traits and diseases (Bulik-Sullivan et al, 2015b), all using GWAS summary-level results data (Table 1).

The chief limitation of using LD score regression to estimate genetic correlations to date has been a practical one. Publicly available GWAS meta-analysis results are available from a number of different repositories on the Internet. It is time consuming to locate and download all of these resources for use, particularly as these databases become more numerous. What’s more, each summary results file typically involves different file formats and conventions making data preparation a time consuming exercise. In addition, many GWAS meta-analyses are not made publicly available, requiring the user to proactively invite the relevant investigators to share their results, which also takes a significant amount of time.

Here we describe a centralized database and web interface, LD Hub, which automates the LD score regression analysis pipeline using publically available GWAS summary-level data of individuals with European ancestry. Users of our web-based tool only need to upload summary results for their trait(s) of interest; and the web server will automatically test their results against GWAS results from (currently) 177 other traits/diseases. The proposed database and web interface calculates the SNP heritability for the uploaded phenotype(s), and a genetic correlation matrix across traits. LD Hub allows the user to conduct the analysis on specific phenotypes only or perform a hypothesis free screen across all traits in the database (Table 1). Users have the option of uploading their own results files and the option of adding their GWAS results to the database for inclusion in future releases. The resource is continuously updated and curated every month to include new results from users and publicly available sources alike. The pre-computed genetic correlation matrix will be provided on LD-Hub for all traits included in the database.

## 2 Methods

As summarized in Figure 1, LD Hub includes: 1) Lookup Center: a facility to perform lookups of existing LD score regression results; 2) Database: a GWAS summary-level statistics database, 3) Test Center: a web interface that automates the LD score regression analysis pipeline including the calculation of SNP heritability and genetic correlations, and 4) GWAShare Center: a user contribution and data sharing platform

**Fig 1.**
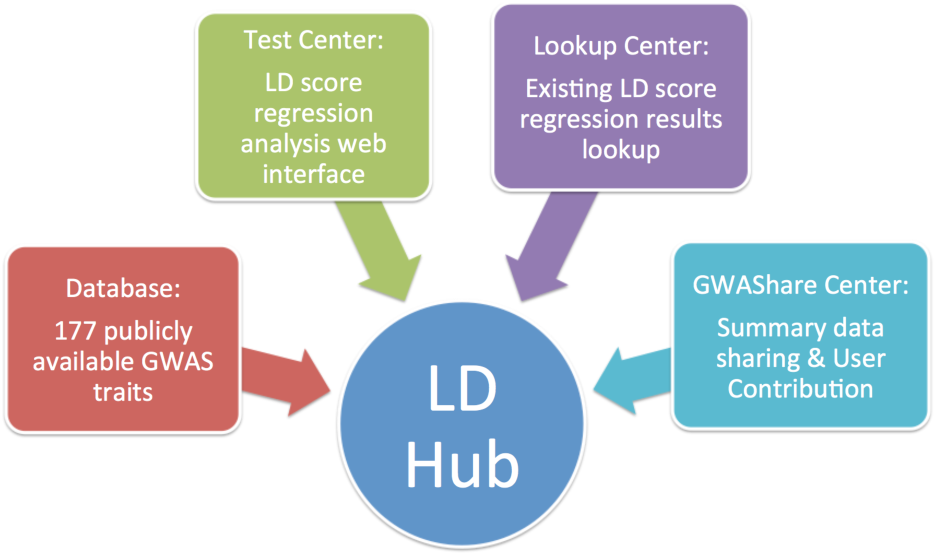
Scope of LD Hub.

### 2.1 LD Hub database

#### 2.1.1 GWAS summary-level data

We cleaned and harmonized 844 publicly available GWAS summary-level data sets from 35 consortia, which included 82 diseases, 154 complex traits, 454 metabolites and 151 immune markers (Hemani et al, 2016). From this database pool, we chose datasets that fit the following selection criteria:

1. Non-sex-stratified
2. Meta-analyses of European populations.
3. Meta-analyses using a GWAS backbone chip only (i.e. exclude meta-analyses involving immuno | metabo | psych | exome chip or GWAS + custom chip)
4. Number of SNPs is large (N>450,000)
5. Number of individuals is large (N>5,000)
6. Mean Chi-square of the test statistics is larger than 1

As shown in Figure 2, after filtering on the selection criteria, genome-wide results for 177 traits were included in LD Hub, of which 23 are GWAS of diseases, 47 are medically relevant risk factors/traits and 107 are metabolites. Table S1, displays descriptive information for each of the GWAS in LD Hub, including, trait name, consortium name, ethnicity, gender, number of cases and controls, sample size, PubMed ID, year of publication, and other relevant information. We used an inclusive strategy for data selection, so we show a list of traits that may return null results in Table S2 due to low Z score of heritability estimate.

**Fig 2.**
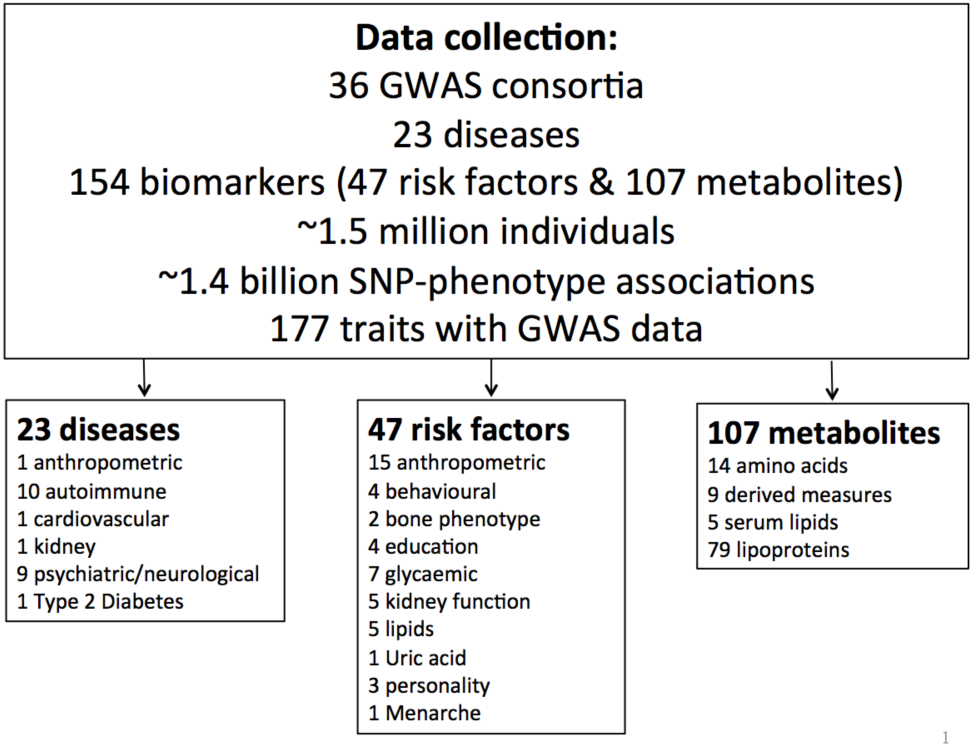
In total, data for 177 traits are included in LD Hub, which consist of 23 diseases, 47 complex traits and 107 metabolites.

#### 2.1.2 LD score information

We pre-calculated LD scores for each SNP using individuals of European ancestry from the 1000 Genomes project (1000 Genomes Project Consortium, 2012). These LD scores are suitable for standard LD score analyses in European populations (i.e. the LD score regression intercept, heritability, genetic correlation, cross-sex genetic correlation).

### 2.2 LD Hub web interface

The LD Hub web interface framework was developed using Python Django as the LD score regression program is also written using Python.

#### 2.2.1 Test Center

The LD Hub web interface provides an automatic LD score regression analysis pipeline for users. As shown in Figure 3, the LD Hub analysis pipeline consists of 5 major steps:

**Fig 3.**
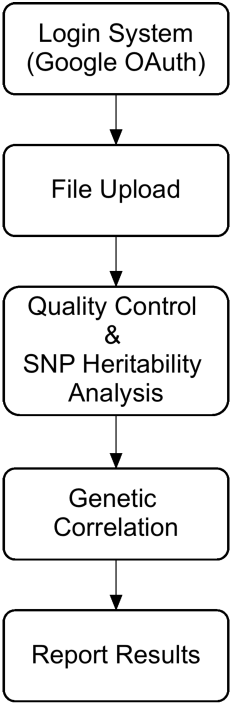
Schematic of LD Hub workflow.

1. User login system: using a Google OAuth
2. File upload system: To run the LD score analysis pipeline, LD Hub requires upload of a file containing summary results data. In the web interface, we provide an example GWAS results file to illustrate the file format required for successful upload and analysis by LD Hub. To save uploading time, each results file should be a white space zipped file in which each row contains the results from a single SNP whilst the columns comprise the following fields:
  a. SNP ID (rs number)
  b. Effect allele of the SNP
  c. Alternate allele of the SNP
  d. Sample size of each SNP (can use an overall sample size if sample size for some SNPs is missing)
  e. A signed summary statistic where the sign refers to the addition of the effect allele (i.e. any statistic that can be converted into a Z-score)
  f. P value of the SNP
  g. Minor allele frequency of each SNP (optional) h) SNP Imputation quality (optional)
3. Quality control and heritability analysis: To standardize the input file, quality control is automatically performed on the uploaded file.
  a. For studies that provide sample MAF, a filter to include SNPs with MAF above 1%.
  b. In order to restrict the analysis to well-imputed SNPs, we filter the uploaded SNPs to HapMap3 SNPs (International HapMap 3 Consortium et al, 2010) with 1000 Genomes EUR MAF above 5%, which tend to be well-imputed in most studies.
  c. If sample size varies from SNP to SNP, remove SNPs with an effective sample size less than 0.67 times the 90th percentile of sample size.
  d. Remove INDELs and structural variants.
  e. Remove strand-ambiguous SNPs.
  f. Remove SNPs whose alleles do not match those in the 1000 Genomes data.
  g. Remove SNPs within the major histocompatibility complex (MHC) region since these often display extreme LD and effect sizes.
  h. Because outliers can unduly influence the regression, we also removed SNPs with extremely large effect sizes (X_1_^2^ > 80).
  The second part of this step is the SNP heritability analysis. The results of this analysis provide a useful indication of whether genetic correlation analysis is likely to be informative (Bulik-Sullivan et al. 2015).
4. Genetic correlation analysis. If the uploaded GWAS results are likely to provide good statistical power (i.e. heritability H^2^ Z score of > 4), then the LD Hub pipeline will perform genetic correlation analysis on the uploaded GWAS results. Users have the option of selecting which traits they want to include in the analysis.
5. Reporting of results.

#### 2.2.2 Lookup Center

Another feature of the LD Hub web interface is the heritability and genetic correlation ‘lookup’ function for existing GWAS results currently in the database. In the current version (v1.0), we provide SNP heritability and genetic correlation results. A forest plot for heritability and a genetic correlation matrix plot are also provided.

#### 2.2.3 GWAShare Center

We aim to promote sharing of summary GWAS results data. We encourage users of LD Hub to upload their GWAS results for curation into the database. We will update the database regularly and allow other users to use the shared data for LD score regression analyses, which will then benefit the whole human genetics community.

### 2.3 Case study of Atopic dermatitis

In order to illustrate the utility of LD Hub, we conduct an analysis using summary results data from a large GWAS of atopic dermatitis (AD) for 40835 (10,788 cases and 30,047 controls, sample prevalence: 0.264) European individuals (i.e. the discovery cohorts from this paper minus results from 23ANDME) (EAGLE consortium 2015). In total, 11059640 SNPs were included in this meta-analysis. Since AD is influenced by a gene of major effect (i.e. filaggrin) which could bias estimates from LDHub, we excluded this region from the uploaded results file. After quality control, 1215002 SNPs were selected for upload.

**Fig 4.**
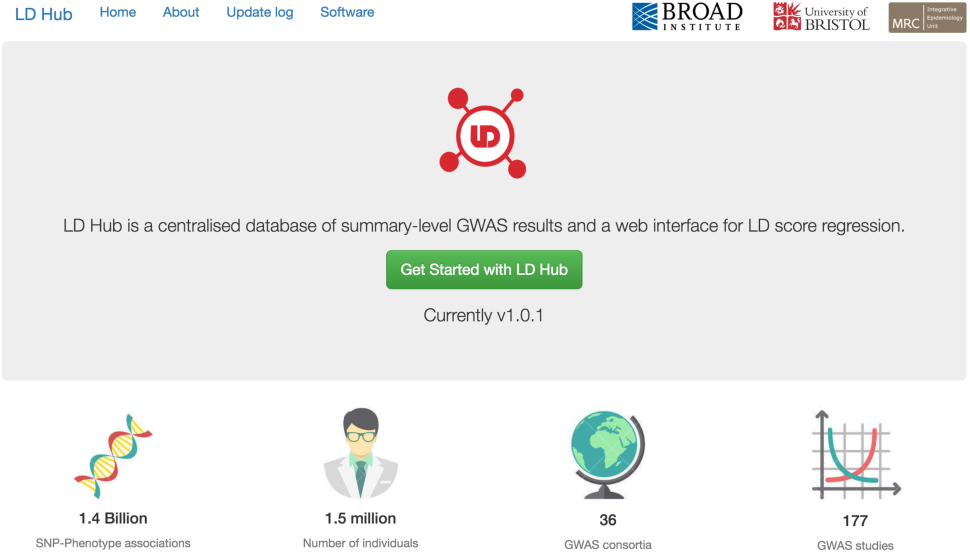
Screen shot of LD Hub web interface.

## 3 Results

### 3.1 Validation of LD Hub analysis results

We tested the validity and functionality of LD Hub by replicating previously reported results from the original LD Score regression suite of papers (Bulik-Sullivan et al, 2015a, Bulik-Sullivan et al, 2015b).

We compared SNP heritability results between LD Hub and previously reported LD score regression results (Bulik-Sullivan et al, 2015a). As shown in Table S3, the Mean *χ*^2^, *λ*_GC_ and Intercept results are almost the same. The minor discrepancy is a consequence of using slightly different quality control processes for LD Hub than what was used in the original LD Score regression paper. Results for SNP heritability of 177 traits are shown in Table S1.

We also compared the genetic correlation analysis results between LD Hub and previously reported results (Bulik-Sullivan et al, 2015b). As shown in Figure 5, the genetic correlation and standard error of genetic correlation estimates are consistent with previously reported LD score regression genetic correlation results. A comparison of the genetic correlation results of (previously reported) 49 traits is shown in Table S4.

**Fig 5.**
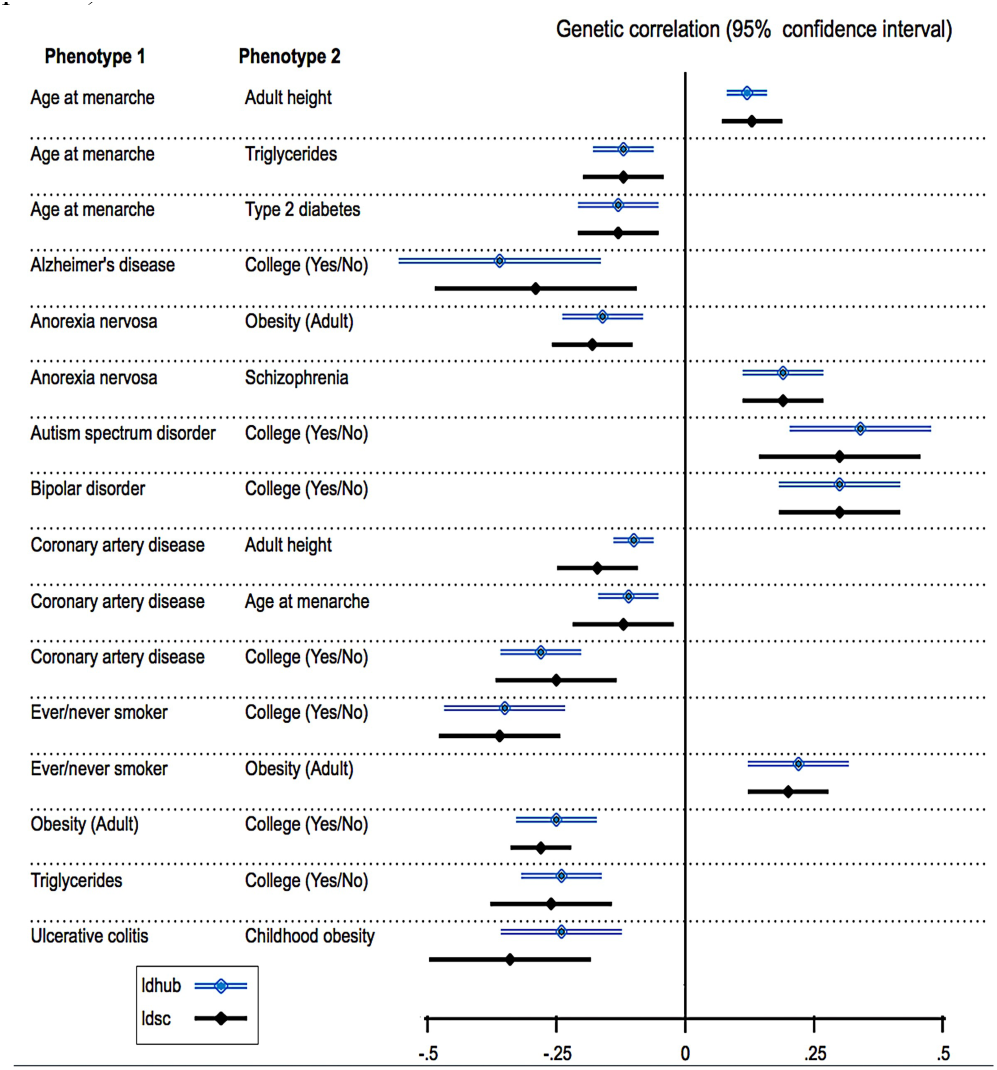
Comparison of genetic correlation results between LD Hub and previously reported LD score regression results.

Double blue lines represent genetic correlation results from LD Hub;and the black single lines represent genetic correlation results from previously reported LD score regression results. The discrepancies can be attributed to the minor changes in the quality control processes and the replacement of some GWAS results with the more recent versions.

### 3.2 Case study: Atopic Dermatitis

Table 2 shows the SNP heritability for AD. The figure of 7.8% is low particularly compared to the heritability estimates from twin studies of eczema where figures exceeding 80% are not uncommon (Bataille et al. 2012). This could be for a number of reasons including the fact that genomic control correction in the individual meta-analysis studies causes downward bias, the filaggrin regions of the genome were excluded from the analysis, and the fact that LD score regression provides an estimate of the overall proportion of additive genetic variance tagged by SNPs in the GWAS panel (i.e. SNP heritability), rather than total heritability *per se*. However the greatest contributing factor is likely to be the case definition of AD used in the EAGLE consortium paper which is extremely heterogeneous, relying often on self-report or retrospective recall which will introduce substantial measurement error into the analysis (and hence decrease heritability estimates). Our results strongly suggest that reanalysis using a more precise definition of eczema would result in a cleaner phenotype and consequently increase the number of genome-wide significant loci detected.

**Table 2.** SNP heritability for atopic dermatitis. H^2^ and SE_H^2^ refer to the SNP heritability and standard error of the SNP heritability.

| Type of Heritability Scale | $H^2$ | $SE\_H^2$ | $\lambda_{GC}$ | Mean $\chi^2$ | Intercept |
| --- | --- | --- | --- | --- | --- |
| Observed Scale | 0.071 | 0.016 | 1.053 | 1.080 | 1.034 |
| Liability Scale | 0.078 | 0.018 | 1.053 | 1.080 | 1.034 |

Table 3 displays estimated genetic correlations between AD and several immune mediated diseases and lung cancer recorded in LD Hub. As expected, the estimated genetic correlation (rG) between AD and asthma was strongly significant and positive. We also note that the rG between AD and Crohn’s disease was moderate, significant and positive, perhaps reflecting substantial overlap between currently known loci for both conditions (Paternoster et al. 2015). rG did not differ significantly from zero for the other traits, although the point estimates for several were moderate indicating that follow up when larger samples become available may be justified.

**Table 3.**
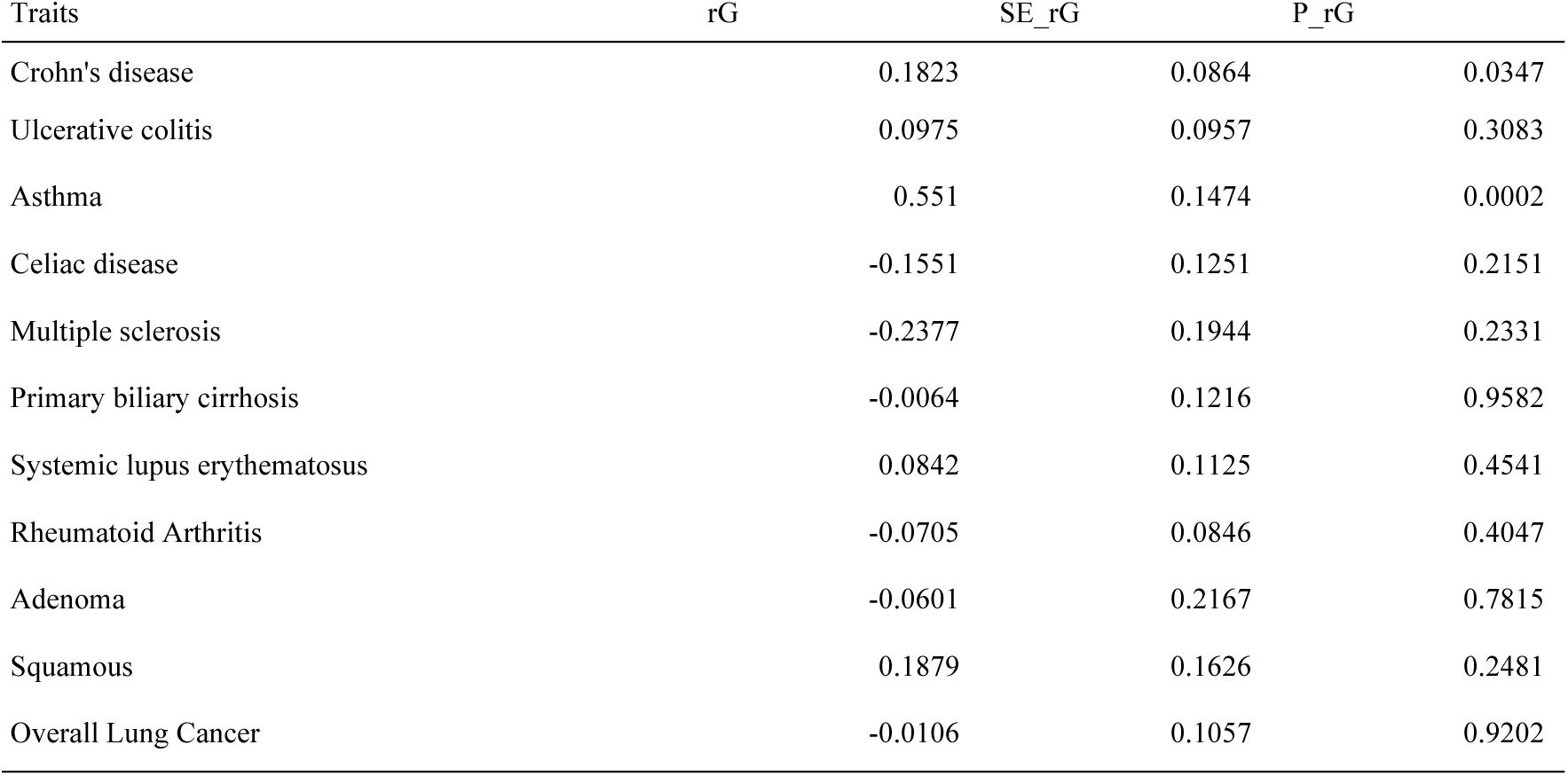
Genetic correlation between atopic dermatitis and other immune mediated diseases. rG refers to the genetic correlation between two traits, SE_rG is the standard error of the genetic correlation, P_rG is the p value of the genetic correlation. The full set of genetic correlation results between AD and other selected traits are included in Table S5.

## 4 Discussion

In this paper, we describe LD Hub (accessible at http://ldsc.broadinstitute.org/), a web-based utility that centralizes and harmonizes summary-level GWAS results data, and automates LD Score regression analysis (Bulik-Sullivan et al, 2015a, Bulik-Sullivan et al, 2015b).

GWAS meta-analysis summary statistics are increasingly being made publicly available. Our database (currently) utilizes results from 177 different GWAS, which includes all publicly available GWAS summary results suitable for LD Score regression (Bulik-Sullivan et al., 2015a). However, this represents a small proportion of the traits represented in the GWAS Catalog (https://www.ebi.ac.uk/gwas/) (Hindorff et al; Welter et al, 2014). There is thus an urgent need for increased sharing of GWAS meta-analysis results in order to realize the full potential of techniques that utilize summary results data such as LD score regression. LD Hub provides a natural platform for the distribution of summary results data that can be utilized by the whole genetics community.

There are four major advantages of using our database and web interface:

1. Users of LD Score regression currently spend most of their time reformatting, harmonizing and managing summary results data rather than running the ‘actual’ analyses. LD Hub minimizes the proportion of time spent on the former so that users can focus their attention on interpreting interesting genetic correlations and SNP heritabilities.
2. Users who do not have a computational background will find the interface easier to use
3. The software is computationally very fast. The current version (v1.0) can return the systematic analysis results to the user within few hours. A queuing system has been introduced to prevent the server from crashing.
4. As users upload and share their own summary GWAS results, the resource becomes increasingly useful. We envisage LD Hub as a useful hypothesis generating tool, providing an easy method of screening hundreds/thousands of traits for interesting genetic correlations that could subsequently be followed up in further detail by other approaches such as pathway analysis (Segre et al. 2010) or Mendelian randomization (Davey-Smith & Ebrahim 2003). For example, under most models, a causal relationship between two heritable traits should induce a genetic correlation between the two phenotypes (assuming individual differences in the causal trait are influenced by genetic variation). LD Hub could be used to screen a large number of putatively causally related phenotypes quickly and easily for evidence of genetic correlation, and the most promising candidate pairs could then be followed up by selecting appropriate genetic instruments and performing formal instrumental variables analysis (Evans & Davey-Smith, 2015, Hemani et al, 2016). This framework could be particularly useful in the dissection of high dimensional molecular networks where the number of possible pair-wise relationships may be extremely large. For LD Hub, we list few suggestions / limitations here:

1. In order for estimates of the genetic correlation to be reliable we suggest that traits uploaded meet the following criteria
  - Heritability (H^2^) Z score is at least > 1.5 (optimal > 4)
  - Mean Chi square of the test statistics > 1.02
  - The intercept estimated from the SNP heritability analysis is between 0.9 to 1.1
2. As we aim to provide an analysis pipeline that is as systematic as possible, we used a very inclusive strategy for data selection, where we expect a very small proportion of the analyses (especially for the traits listed in Table S2) to return null results.
3. LD Hub is currently designed for GWAS studies involving European populations exclusively. As the number of publicly available GWAS involving other ethnicities increases we will extend LD Hub to include these. In summary, due to the growing availability of summary-level data, our database together with the web interface will maximize the potential of GWAS summary-level data for heritability and genetic correlation analyses.

## Acknowledgements

We would like to thank Kaitlin Wade, Vanessa Tan, Ryan Langdon and James Yarmolinsky for helping curate the GWAS data information.

